# Potent and specific human monoclonal antibodies against SARS-CoV-2 Omicron variant by rapid mRNA immunization of humanized mice

**DOI:** 10.1101/2022.03.17.484817

**Authors:** Ping Ren, Lei Peng, Zhenhao Fang, Kazushi Suzuki, Paul Renauer, Qianqian Lin, Meizhu Bai, Luojia Yang, Tongqing Li, Paul Clark, Daryl Klein, Sidi Chen

**Author notes:** Correspondence: SC. Co-first authors (Ping Ren, Lei Peng). Lead contact: SC, (lab).

## Abstract

The Omicron variant (B.1.1.529) of SARS-CoV-2 rapidly becomes dominant globally. Its extensive mutations confer severe efficacy reduction to most of existing antibodies or vaccines. Here, we developed **RAMIHM**, a highly efficient strategy to generate fully human monoclonal antibodies (mAbs), directly applied it with Omicron-mRNA immunization, and isolated three potent and specific clones against Omicron. Rapid mRNA immunization elicited strong anti-Omicron antibody response in humanized mice, along with broader anti-coronavirus activity. Customized single cell BCR sequencing mapped the clonal repertoires. Top-ranked clones collectively from peripheral blood, plasma B and memory B cell populations showed high rate of Omicron-specificity (93.3%) from RAMIHM-scBCRseq. Clone-screening identified three highly potent neutralizing antibodies that have low nanomolar affinity for Omicron RBD, and low ng/mL level IC50 in neutralization, more potent than majority of currently approved or authorized clinical RBD-targeting mAbs. These lead mAbs are fully human and ready for downstream IND-enabling and/or translational studies.

## Introduction

SARS-CoV-2 has rapidly spread across the world, causing a global pandemic of coronavirus disease 2019 (COVID-19) and posing a serious threat to global healthcare systems(Corbett et al., 2020; Lu et al., 2020; Zhou et al., 2020). To date, SARS-CoV-2 has infected hundreds of millions of people and caused millions of deaths worldwide(Dejnirattisai et al., 2022b). Although multiple approved vaccines and neutralizing mAbs were rapidly deployed(Baden et al., 2021; Krammer, 2020; Polack et al., 2020; Sadoff et al., 2021; Weinreich et al., 2021), the emergence of new variants with mutations in spike (S) glycoprotein that could escape the antibody response further threaten the protective immune responses from infection, vaccination or antibody therapies (Dhar et al., 2021; Faria et al., 2021; Tegally et al., 2021).

Recently, the B.1.1.529 variant of SARS-CoV-2 was declared as variant of concern (VoC) and designated as Omicron by the World Health Organization (WHO)(Karim and Karim, 2021; Scott et al., 2021). Compared with previous VOCs, the Omicron variant is particularly concerning due to a high number of mutations, especially in the spike protein relative to the ancestral virus of SARS-CoV-2. Notably, 15 Omicron mutations were distributed at the receptor-binding domain (RBD), including G339D, S371L, S373P, S375F, K417N, N440K, G446S, S477N, T478K, E484A, Q493R, G496S, Q498R, N501Y, and Y505H, which is the primary target of serum neutralizing antibodies elicited by infections or vaccines(Piccoli et al., 2020). Among the mutations and indels in N-terminal domain (NTD), the 143-145 deletion is located in the antigenic supersite targeted by most of NTD neutralizing antibodies and is predicted to mediate immune escape of NTD-targeting antibodies(Cao et al., 2021; Cerutti et al., 2021; McCallum et al., 2021). SARS-CoV-2 spike contains a unique S1/S2 furin cleavage site (681-685aa in WT, PRRAR), which is associated with SARS-CoV-2 transmissibility(Johnson et al., 2021) and pathogenesis(Johnson et al., 2021), and is not present in other group 2B coronaviruses. Two Omicron mutations (N679K and P681H) adjacent to the furin site add additional positive charged residues to this short basic stretch and are predicted to enhance transmissibility. A series of recent studies have shown that the mutations in Omicron variant lead to marked reduction of neutralizing activity from vaccination and from approved or emergency-authorized therapeutic monoclonal antibodies (mAbs) (Callaway, 2021; Cao et al., 2021; Cele et al., 2021b; Ju et al., 2020; Planas et al., 2021). These mutations in Omicron variant renders the vast majority of the originally authorized monoclonal antibodies ineffective (Cao et al., 2021; Hoffmann et al., 2022), causing them to be no longer recommended in the COVID-19 treatment guidelines for patients with Omicron variant infections.

Data on initial epidemiological studies demonstrated the Omicron variant is leading the fourth wave of the SARS-CoV-2 pandemic worldwide, potentially due to its higher transmissibility and immune evasion of SARS-CoV-2 neutralizing antibodies (Cao et al., 2021; Liu et al., 2021; Meo et al., 2021; Starr et al., 2021; VanBlargan et al., 2022; Wolter et al., 2022). Although Omicron variant appeared to be less severe, in the currently largely vaccinated general population, the enormous number of infections still led to large numbers of hospitalizations and deaths daily. Therefore, it is essential to develop next-generation neutralizing mAbs that retain potency and limit SARS-CoV-2 virus transmission when current vaccines and therapeutic antibodies are compromised(Cele et al., 2021a).

In this study, we developed **RA**pid **m**RNA **I**mmunization of **H**umanized **M**ice (**RAMIHM**), an accelerated animal immunization approach for neutralizing mAb discovery. The principle of this approach is to utilize the high doses of antigen-specific LNP-mRNA to frequently immunize immunoglobulin (Ig) humanized mice within 2 weeks, for isolation of high potency neutralizing mAbs against the targeted antigen. We applied this approach directly with Omicron spike-encoding mRNA, used customized single cell BCR sequencing (scBCR-seq) to obtain the human variable region sequences from enriched B cell clonotypes, and generated potent and specific fully human antibodies against the Omicron variant.

## Results

### Development of RApid mRNA Immunization of Humanized Mice (RAMIHM), a highly efficient strategy to identify fully human monoclonal antibodies

To date, two-dose SARS-CoV-2 mRNA-based vaccination strategy has been demonstrated to effectively induce humoral and cellular immunity to SARS-CoV-2, including the ancestral virus (ancestral, reference, wildtype (WT), Wuhan-1, or WA-1, identical sequences), and its VoCs such as Delta variant(Lopez Bernal et al., 2021; Naranbhai et al., 2021). However, a number of recent studies demonstrated that the SARS-CoV-2 Omicron variant has substantial changes in its genome, especially the spike protein (**Fig. 1A**), and illustrated dramatically decreased neutralizing titers in convalescent or vaccinated recipients, causing waning immunity and massive breakthrough infections(Carreno et al., 2021; Cele et al., 2021b; Dejnirattisai et al., 2022b; Hu et al., 2022). Importantly, nearly all antibodies initially developed against the ancestral virus have substantially dropped, or completely lost, the neutralization ability against Omicron(Cao et al., 2021; Dejnirattisai et al., 2022a; Liu et al., 2021; Planas et al., 2021). For example, several of the currently approved or emergency authorized mAbs have their binding interfaces impacted by the Omicron mutations (**Fig. 1B**). Therefore, next-generation neutralizing antibodies are needed in speed. To combat the rapidly evolving VoCs, for example the current resurgence of the Omicron pandemic, it is important to have the ability to rapidly develop translatable human neutralizing mAbs to quickly react to the needs of new therapeutics. We thus developed a novel antibody discovery approach named RAMIHM, with repetitive intramuscular injections using high doses of LNP-mRNA, followed by B cell isolation, antigen enrichment and single B cell sequencing (**Fig. 1C**). We applied this directly with Omicron-spike-encoding LNP-mRNA to induce Omicron-specific immune responses for isolation of Omicron-targeting mAbs.

**Figure 1.**
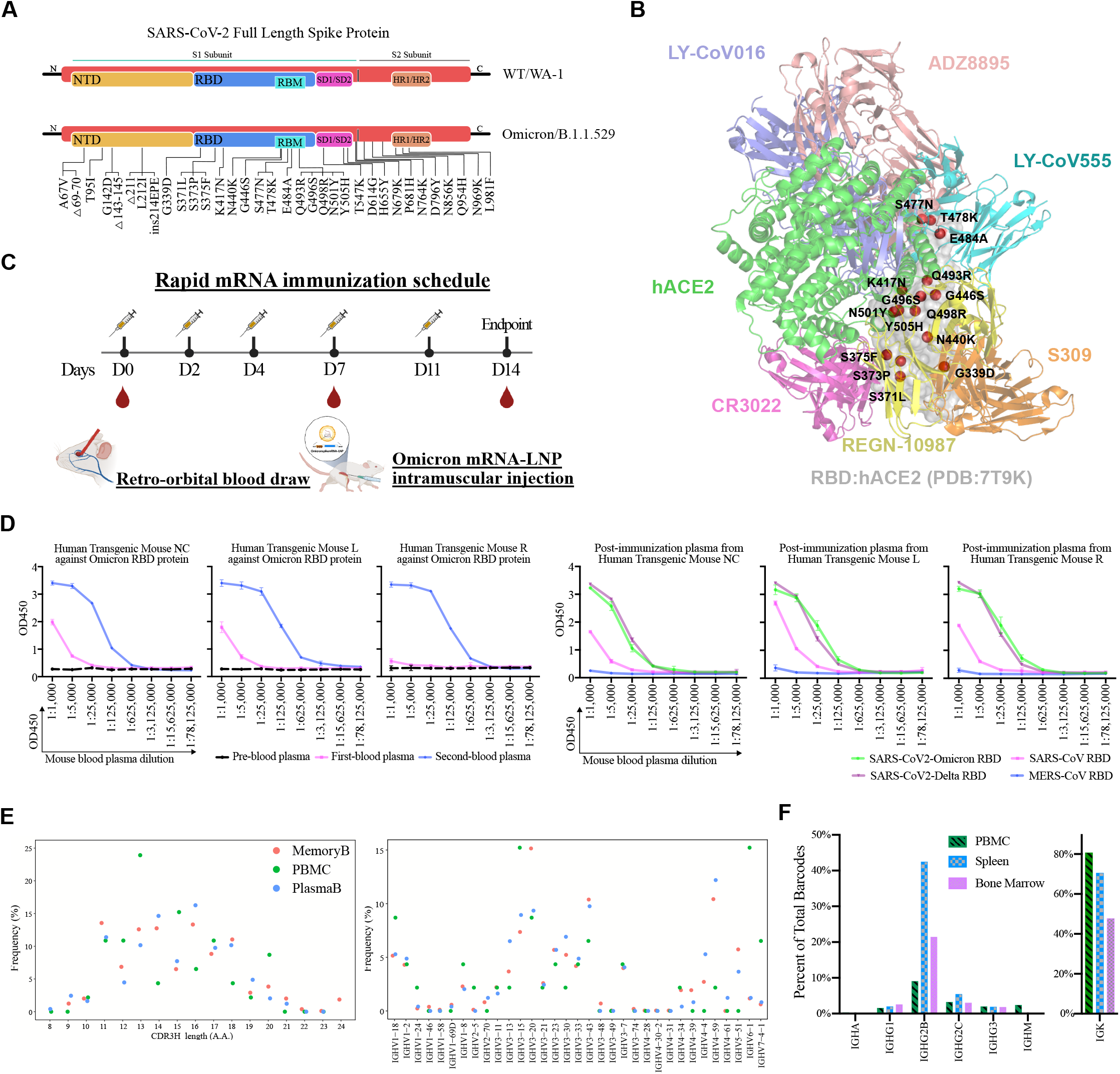
Development of RAMIHM for rapid discovery of fully human monoclonal antibodies and application with Omicron mRNA immunization. **A**, Schematic showing the domain arrangement of the SARS-CoV-2 WT spike and its recent variant SARS-CoV-2 B.1.1.529 (Omicron). Mutations present in Omicron spike protein are labeled. Full-length of Omicron spike gene was synthesized to construct Omicron-specific mRNA-lipid nanoparticle and Omicron-specific pseudo-virus. **B**, Footprint of SARS-CoV-2 RBD-directed antibodies. The SARS-CoV-2 Omicron RBD/hACE2 structure was downloaded from PDB 7T9K, approved or authorized antibodies are labeled. **C**, Schematic illustration of immunization and blood sample collection. Three humanized mice were repetitively immunized with Omicron LNP-mRNA as immunogen. 10μg of Omicron LNP-mRNA were given for each mouse on day0, day2 and day4 and day7, and followed by 20μg of Omicron LNP-mRNA were injected on day11. Retro-orbital blood was collected on day0, day7 and day14. Plasma was isolated from blood for downstream experiments. **D**, Anti-plasma titer determination. Upper panel, all plasma samples were serially 5-fold diluted from 1:1000 and assayed by a direct coating ELISA with Omicron RBD protein coated plate. Error bars represent mean ± SEM of triplicates with individual data points in plots. Lower panel, all post-immunized plasma samples (2^nd^ blood) were serially 5-fold diluted from 1:1000 and assayed by a direct coating ELISA with selected pan-CoV-RBD proteins coated plate, respectively. Error bars represent mean ± SEM of triplicates with individual data points in plots. **E**, B cell characterization by customized scBCR-seq profiling. Left panel, Distribution of heavy chain complementarity-determining region 3 (HCDR3) length in each B cell group (Memory B, Plasma B and PBMC) from Omicron-RAMIHM mice. Right panel, distributions of heavy chain V-segment in each B cell group (Memory B, Plasma B and PBMC) from Omicron-RAMIHM mice. Total number of single cells sequenced with BCRs (Memory B library, n = 2,646; Plasma B library, n = 617; PBMC library, n = 239; Total n = 3,502). **F**, Ig class distributions of Omicron-RAMIHM mice’s clonotypes. Distribution and frequency analysis of immunoglobulin isotypes usage in spleen, bone marrow and PBMC from Omicron-RAMIHM mice. Source data and additional statistics for experiments are in supplemental excel file(s).

Using Omicron-specific LNP-mRNA that contains lipid nanoparticle formulated mRNA encoding the HexaPro engineered full length of Omicron spike glycoprotein (Methods), we first characterized the biophysical integrity of these LNP-mRNAs (**Fig. S1A, S1B**), and validated the expression of functional Omicron spike protein surface expression via human ACE2 (hACE2) staining of LNP-mRNA transfected HEK293 cells (**Fig. S1C**). Next, we performed administration of four 10μg doses and one 20μg dose of Omicron specific-mRNA LNP in 3 IgG-humanized mice, collected retro-orbital blood samples from each humanized mouse before and after booster immunization. Blood samples were labeled as pre-, 1^st^-, or 2^nd^ immunization draw depend on collection sequence (**Fig. 1C**). Antibody titers were measured using serial plasma dilutions on ELISA plates coated with recombinant Omicron RBD protein. Binding activity was visualized using anti-mouse IgG antibodies at 450nm optical density (OD). Three sequential plasma samples showed increasing vaccine-elicited antibody responses during each blood collection (**Fig. 1D**). All post-immunized plasma samples (2^nd^ blood) showed strong reactivity to the recombinant SARS-CoV-2 Omicron RBD protein antigen (**Fig. 1D**). In addition, all these samples also showed strong cross-reactivity to recombinant SARS-CoV-2 Delta RBD protein, and intermediately cross-reactivity to recombinant SARS-CoV RBD protein, but no cross-binding to recombinant MERS-CoV RBD protein (**Fig. 1D**). Together, these results demonstrated that Omicron-specific rapid mRNA immunization (Omicron-RAMIHM) elicited strong anti-Omicron plasma in IgG humanized mice in two weeks, which also contains broader reactive antibodies against other variant and coronavirus species such as SARS-CoV-2 Delta and SARS-CoV.

### Customized single cell BCR sequencing (scBCRseq) mapped the IgG clonal repertoires of Omicron-RAMIHM animals

To obtain SARS-CoV-2 Omicron RBD-reactive B cells, we isolated spleen, lymph nodes, bone marrow and whole blood from Omicron-RAMIHM mouse, and collected three different types of B cells (memory B cell, plasma B cells, and peripheral blood mononuclear cells) by using different isolation procedures (Methods), for B cell repertoire mapping and reactive BCR identification via scBCR-seq. To prepare memory B cells enriched library, we used mouse memory B cell isolation kit to obtain total memory B cells from fresh spleen and lymph nodes, and baited SARS-CoV-2 Omicron RBD specific memory B cells by enrichment using recombinant Omicron-RBD proteins from isolated memory B cell subsets (Memory B library). To generate plasma B cells enriched library, we applied anti-mouse CD138^+^ plasma cell isolation to isolate CD138^+^ plasma B cells from freshly isolated raw bone marrow cells (Plasma B library). To generate peripheral blood mononuclear cells library, we isolated peripheral blood mononuclear cells (PBMCs) by centrifugation using PBMC isolation method from whole blood (PBMC / Peripheral B library). We subjected each single cell BCR sequencing library with input of approximately 10,000 fresh cells from above. After sequencing, we analyzed a total of 3,502 single B cells, and obtained 2,558 paired heavy- and light-chain variable regions of antibody sequences (**Fig. 1E**). To examine the IgG clonal repertoires from the scBCRseq data, we first examined B cell clonotypes, by calculating the frequencies of cells observed for the clonotype and distributions of identical CDR3 region for both heavy and light chains in pairs. By analyzing the BCR repertoires, we mapped the landscape of BCR populations in the Memory B, Plasma B and Peripheral B / PBMC in Omicron-RAMIHM immunized mouse **(Fig. S2, Dataset S1**).

The SARS-CoV-2 Omicron RBD-specific antibodies had a relative enrichment for IGVH3-7, IGVH3-15, IGVH3-20, IGVH3-23, IGVH3-30, IGVH3-33, IGHV3-43, and IGVH4-59, analyzed from 3 individual BCR libraries (**Fig. 1E**). A range of lengths between 8-24 aa was observed for these BCR CDRH3s (**Fig. 1E**). Interestingly, a large portion of IgG2B-expressing B cells were identified from three B cell type isolations (**Fig. 1F**), a signature of potential involvement of Th2 cells in B cells maturation and class switch in these mice undergoing the Omicron-RAMIHM procedure. By analyzing the Ig heavy chain (IGH) and light chain (IGK) paring, we also mapped out the overall, enriched and the top 10 heavy-and light-chain V/J segment recombination in these B cell populations (**Fig. 2A-B, Fig. S3, Dataset S1**). In summary, scBCRseq data mapped the clonal repertoires and revealed enriched IgG clonotypes in the peripheral blood, plasma B cell and memory B cell populations in Omicron-RAMIHM humanized mouse.

**Figure 2.**
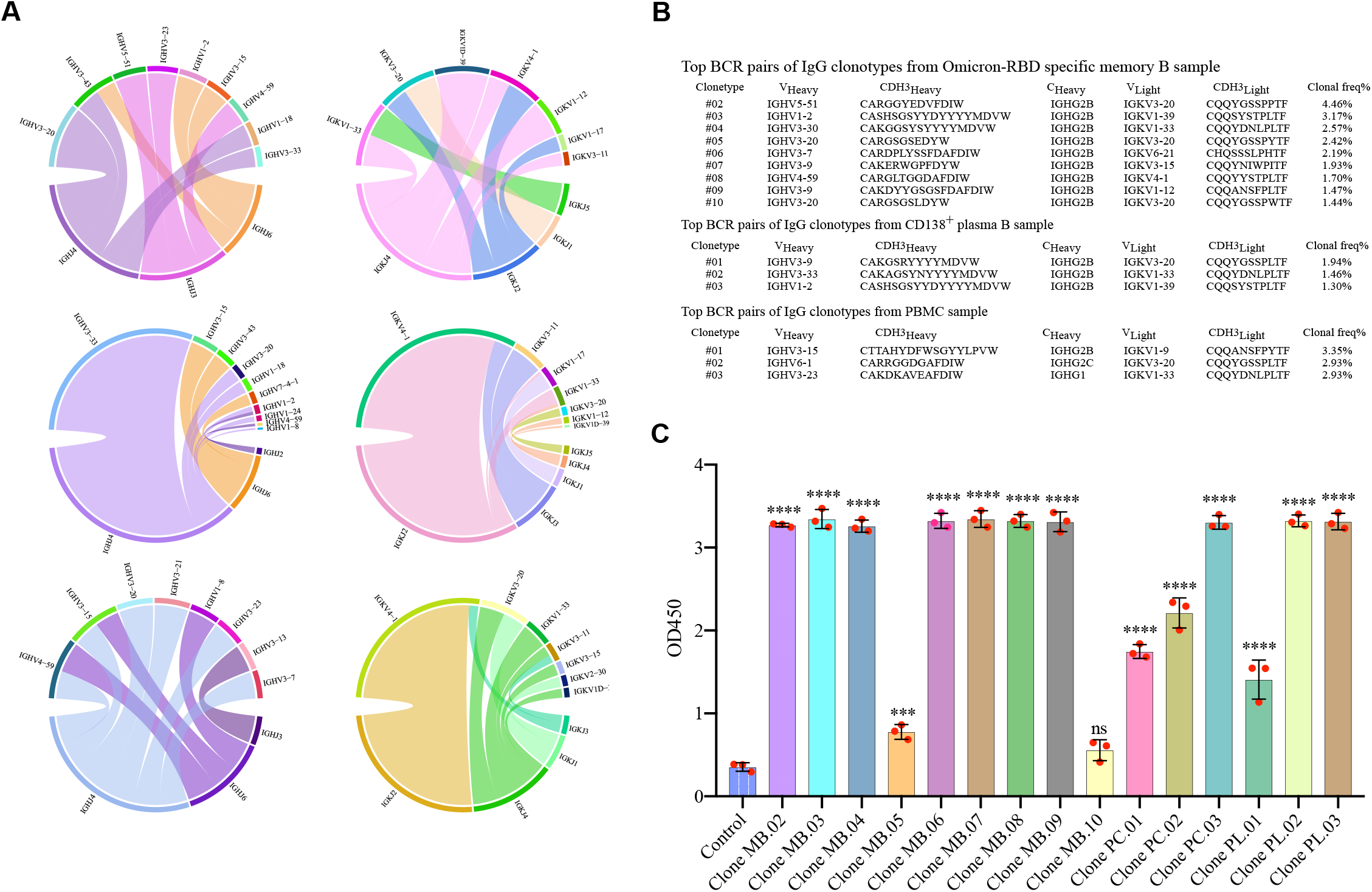
Identification of fully human Omicron-specific monoclonal antibodies against Omicron. **A**, Distribution of top10 heavy- and light-chain V/J segment recombination. Chord diagrams (circos plots) showing the distribution of top10 heavy- and light-chain V and J gene-segment recombination obtained in each representative library. Interconnecting lines indicate the relationship between antibodies that share V and J gene-segment at both IGH and IGL. Top to bottom: Memory B library, PBMC library, and Plasma B library. **B**, Single B cell variable chains for antibody cloning. Variable (V) genes and CDR3 lengths for paired heavy- and light-chains of top-enriched clones to SARS-CoV-2 Omicron from single BCR sequencing. **C**, ELISA of mAbs supernatant binding specificity against Omicron RBD protein. All full length mAb clones from single BCR sequencing and control were evaluated against Omicron RBD protein coated on the ELISA plate and binding activity was recorded at an optical density (OD) of 450nm. Triplicate datapoints (n = 3 each). In this figure: Data are shown as mean ± s.e.m. plus individual data points in dot plots.Statistics: One-way ANOVA was used to assess statistical significance. Each mAb clone was compared to control. Multiple testing correction was made to correct the p values. Two-sided tests were performed. The p-values are indicated in the plots. Statistical significance labels: * p < 0.05; ** p < 0.01; *** p < 0.001; **** p < 0.0001. Non-significant comparisons are not shown, unless otherwise noted as n.s., not significant. Source data and additional statistics for experiments are in supplemental excel file(s).

### Identification of Omicron-specific functional mAb clones from top-ranked paired human Ig chains of Omicron-RAMIHM animals

To test whether the most enriched BCRs in these B cell populations are Omicron-reactive, we selected a panel of BCRs for recombinant mAb expression, including 3 from peripheral blood, 3 from plasma B and 9 from memory B cell populations (**Fig. 2C**). In order to functionally analyze the antibody response to SARS-CoV-2 Omicron RBD, we cloned paired heavy- and light-variable segments into human IgG1 expression vectors (**Fig. 3A**), and used the Expi293F mammalian expression system to produce selected mAbs. Thereafter, we used SARS-CoV-2 Omicron RBD-specific ELISA to determine antibody binding by using transfected culture supernatants that contain secreted antibodies. As a result, almost all of the top-enriched antibody clones collectively from peripheral blood, plasma B cell and memory B cell populations are reactive to Omicron RBD (14/15 reactive, 1/15 slightly reactive), showing a high rate of antigen-specificity (14/15, 93.3%) from Omicron-RAMIHM-scBCRseq (**Fig. 2C**). Ten out of fifteen (10/15) selected clones showed potent binding capacity against recombinant SARS-CoV-2 Omicron RBD proteins, 4/15 showed moderate binding, an 1/15 showed relatively weak binding (**Fig. 2C**). These results indicated that RAMIHM is a highly effective approach for generating and isolating antigen-specific mAbs.

**Figure 3.**
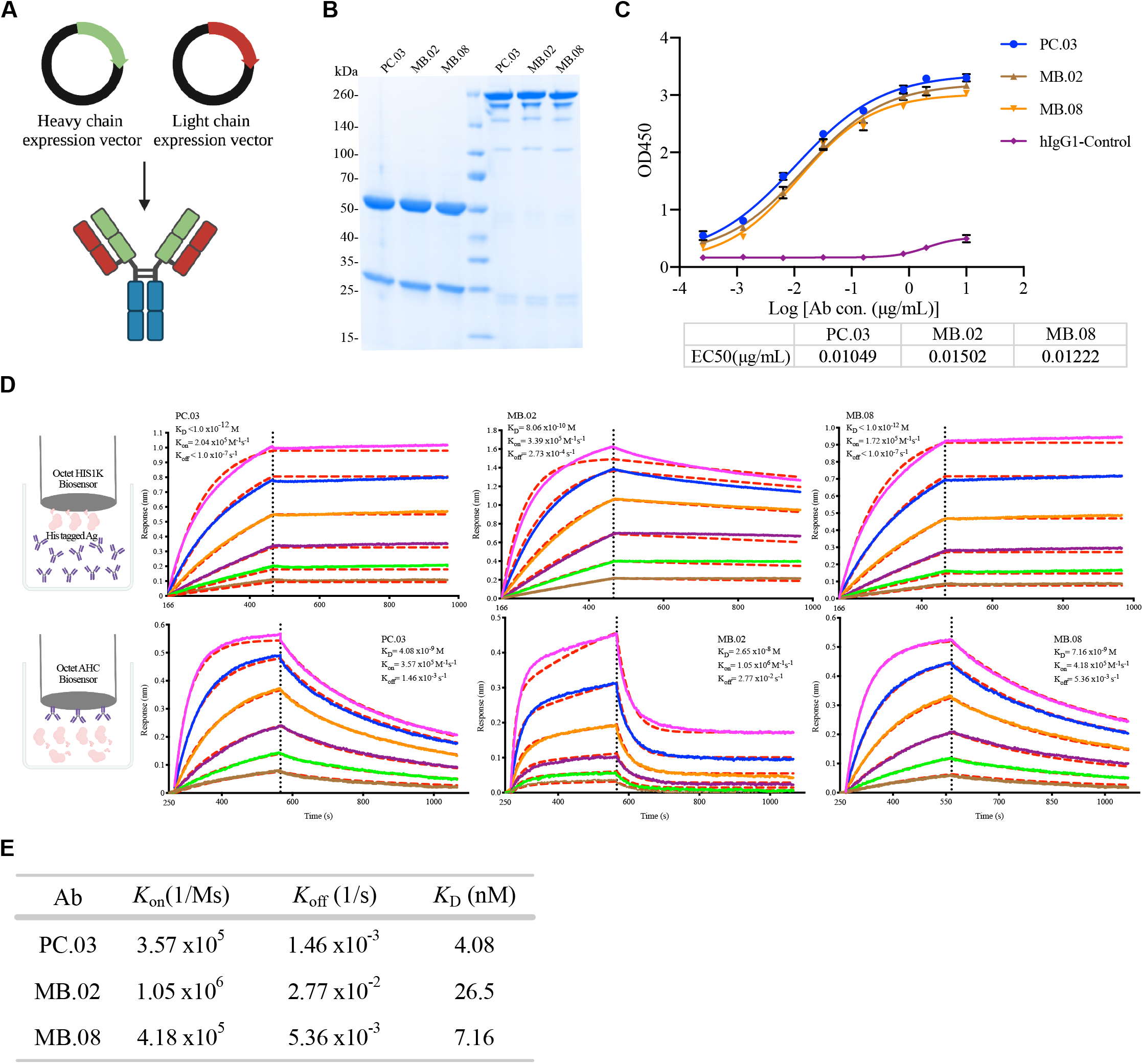
Biophysical and functional characterization of lead clones of Omicron-specific antibodies. **A**, Schematic of human IgG1 mAb production. **B**, SDS-PAGE analysis of purified mAbs under nonreducing and reducing (10mM DTT) conditions. Four micrograms of purified protein were analyzed using a Novex WedgeWell 4-20% (wt/vol) Tirs-Glycine gel. **C**, Graph shows leading Omicron mAbs reactivity. The ELISA EC50 values were calculated by Prism V8.0 software using a four-parameter logistic curve fitting approach. Error bars represent mean ± SEM of triplicates with individual data points in plots. **D**, Binding characteristics of the neutralizing mAbs determined by using BLI. Upper panel, recombinant SARS-CoV-2 Omicron RBD were covalently immobilized onto a HIS1K sensor, all measurements were performed by using a serial 2-fold dilution of purified mAbs, starting from 50nM (Magenta) to 1.56nM (Brown). Lower panel, purified mAbs were immobilized onto an AHC sensor, all measurements were performed by using a serial 2-fold dilution of soluble SARS-CoV-2 Omicron RBD, starting from 50nM (Magenta) to 1.56nM (Brown). Global fit curves are shown as red dashed lines, The vertical black dotted dashed lines indicate the transition between association and disassociation phases. **E**, Summary data of BLI (D) results. Source data and additional statistics for experiments are in supplemental excel file(s).

To further screen for highly potent functional mAbs, we recombinantly expressed these 15 mAb candidate clones in mammalian system and tested their neutralization ability against the Omicron variant. By screening the mAbs from culture supernatants by neutralizing assay using a spike-based SARS-CoV-2 Omicron pseudovirus system, we found 3 clones with obvious neutralization activity against Omicron pseudovirus (**Fig. S4A-B**). We chose these top 3 clones (named as PC.03, MB.02, and MB.08) for further development and characterization.

### Characterization of fully human lead clones with strong binding to Omicron RBD

We purified the three leading clones, PC.03, MB.02, and MB.08, by affinity chromatography using Protein A beads and examined antibody purity by SDS-PAGE (**Fig. 3B**). Thereafter, purified leading mAbs were tested for SARS-CoV-2 Omicron RBD reactivity by ELISA and monitored real-time association and dissociation to recombinant SARS-CoV-2 Omicron RBD proteins using the Octet system. The ELISA titration result of lead mAb clones vs. recombinant SARS-CoV-2 Omicron RBD proteins showed that these three mAb clones have EC50s at the level of ∼0.01 µg/mL, suggesting that these mAbs can indeed tightly bind to Omicron RBD (EC50<16ng/mL for all 3 clones) (**Fig. 3C**). Octet results with his-tag Omicron RBD antigen immobilization showed ultra-strong binding (K_*D*_ at 0.8nM for MB.02, and K_*D*_ <1pM for PC.03 and MB.08) (**Fig. 3D**). Noted that this might be contributed by avidity effect due to multi-valent binding, we also performed the reverse Octet assay with antibody immobilization, which measured the single-mAb binding affinity (**Fig. 3D**), and showed that the affinity between these clones to Omicron RBD are at the level of low nanomolar range (**Fig. 3D**). These K_*D*_ values (**Fig. 3E**) showed that the binding strengths of the 3 lead mAbs are stronger than that of hACE2 with Omicron RBD (31.4±11.62nM) (Han et al., 2022). Noted that most of approved or EUA mAbs have much weaker binding with Omicron RBD (Cameroni et al., 2021; McCallum et al., 2022) (summarized in **Table S1**).

To further determine whether these leading mAbs compete for similar epitopes, we performed epitope binning experiments by Octet using an in-tandem assay (**Fig. S5A**). The results have exhibited that PC.03, MB.02, and MB.08 likely share overlapping epitopes (**Fig. S5B-C**). We next measured antibody competition with ACE2, which was quantified as reduction in ACE2 and RBD binding. Consistent with binding affinity findings, these three leading clones showed competitive binding with ACE2 against Omicron RBD (**Fig. S6A-D**).

### Further characterization of fully human lead neutralization mAb clones against Omicron

We then performed neutralization assays for the 3 lead mAbs in purified form, along with other mAbs. We previously identified and developed several potent and specific mAbs against the ancestral virus and the Delta variant, namely clones 2, 6 and 13A(Peng et al., 2021). In a pseudovirus neutralization assay, we found that while clones 2 and 13A can still neutralize Omicron variant, the potency is significantly reduced (by 1-2 orders of magnitude in terms of IC50 values, at 0.396 and 1.761 µg/mL for clone 2 and 13A, respectively) (**Fig. 4A**), a phenomenon similar to other mAbs developed against the ancestral spike(Dejnirattisai et al., 2022a; Liu et al., 2021). In contrast, all three clones, PC.03, MB.02, and MB.08, potently neutralized the Omicron variant, with IC50 values at 0.15 µg/mL (PC.03), 0.09 µg/mL (MB.02), and 0.04 µg/mL (MB.08) (**Fig. 4B; Fig. S7A**). The neutralization potency of the 3 lead Omicron-specific mAb clones are much stronger than those of our prior mAbs and those under prior regulatory approval or EUAs (**Fig. 4, Table S1**). These 3 mAbs however showed no neutralization against the Delta variant (**Fig. S7B**), further suggesting that they are Omicron-specific.

**Figure 4.**
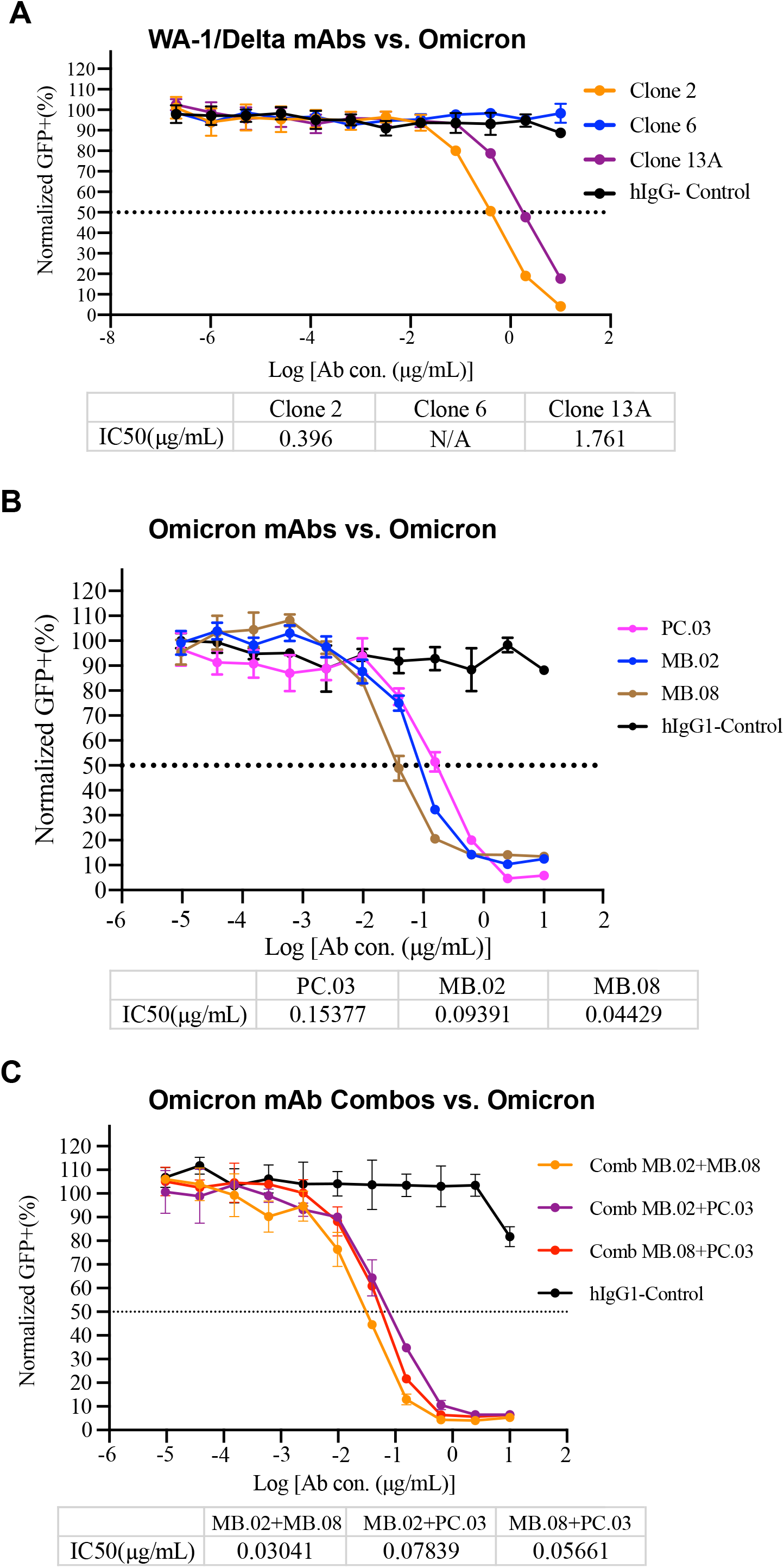
Additional functional characterization of lead clones of Omicron-specific antibodies. **A**, Neutralization assay of SARS-CoV-2 Omicron pseudovirus by WA-1/Delta mAbs. Graph shows the normalized relative GFP signals for detection of 293T cells expressing hACE2, 24h after infection with SARS-CoV-2 Omicron pseudovirus, in the presence of increasing concentration of indicated WA-1/Delta mAbs. **B**, Neutralization assay of SARS-CoV-2 Omicron pseudovirus by leading Omicron mAbs. Graph shows the normalized relative GFP signals for detection of 293T cells expressing hACE2, 24h after infection with SARS-CoV-2 Omicron pseudovirus, in the presence of increasing concentration of indicated Omicron mAbs. **C**, Neutralization assay of SARS-CoV-2 Omicron pseudovirus by leading Omicron mAb combinations. Graph shows the normalized relative GFP signals for detection of 293T cells expressing hACE2, 24h after infection with SARS-CoV-2 Omicron pseudovirus, in the presence of increasing concentration of indicated Omicron mAb combinations (MB.02+MB.08, MB.08+PC.03, MB.02+PC.03). The IC50 values were calculated by Prism V8.0 software using a four-parameter logistic curve fitting approach. Dashed line indicated 50% reduction in viral infectivity. Error bars represent mean ± SEM of triplicates with individual data points in plots. Source data and additional statistics for experiments are in supplemental excel file(s).

In order to test if these clones can be used in combination, we again performed neutralization assays by combining two clones. Interestingly, despite epitope overlap, these mAb clones can still enhance each other’s neutralization capacity, with the best combination being an antibody cocktail of MB.02 + MB.08 (IC50 = 0.03 µg/mL) against pseudotyped SARS-CoV-2 Omicron variant (**Fig. 4C**). In summary, these lead neutralizing mAbs showed that they have high affinity vs Omicron RBD, and strong potency in pseudovirus neutralization, which are at least 2 orders of magnitude more potent than existing clinically approved or authorized SARS-CoV-2 mAbs, where their cocktail combinations can also further enhance the neutralization potency (**Table S1**).

## Discussion

To date, the COVID-19 pandemic has entered into a next stage since the emergence of SARS-CoV-2 Omicron variant, which spread globally in recent months due to higher transmission rates and immune escape (Cao et al., 2021; Liu et al., 2021; Planas et al., 2021; Viana et al., 2022; Volz et al., 2021). The Omicron variant harbors 15 mutations were reported in the RBD domain compared with the ancestral Wuhan-1/WA-1 virus, with 9 of these mutations overlap with ACE2 binding footprint, the mediator of host cell entry. In addition, currently approved vaccines (such as BNT162b2, mRNA-1273, and Ad26.COV2.S) are all designed against the original wild-type SARS-CoV-2 (Jackson et al., 2020; Polack et al., 2020; Sadoff et al., 2021). However, it has been shown that neutralizing antibody responses of sera from convalescent or vaccinated individuals was dramatically decreased with increased time post vaccination to against the emerging variant (Flemming, 2022; Hu et al., 2022; Rossler et al., 2022).

The highly mutated Omicron variant has the potential for evasion of binding and neutralization by the majority of clinically neutralizing mAbs (Cao et al., 2021; Dejnirattisai et al., 2022a; Liu et al., 2021; Takashita et al., 2022; VanBlargan et al., 2022). To experimentally validate this assumption, we previously developed and validated 3 high potency neutralizing mAbs against authentic SARS-CoV-2 ancestral virus and Delta variant (Peng et al., 2021). We found that the Omicron variant, harboring substantially more mutation that prior variant, indeed could completely or partially escape neutralization by existing potent SARS-CoV-2 mAbs including approved or emergency authorized clinical antibodies.

To provide countermeasurements quickly to new VoCs such as the Omicron variant, we developed a highly effective animal immunization approach (RAMIHM) with high-throughput customized single cell BCR sequencing. RAMIHM enables us to obtain potent antigen-specific neutralizing mAbs within 3 weeks, offering the opportunity to rapidly respond the potential risks of emerging new viruses or variants. Compared to other approaches, RAMIHM does not reply on human samples and is fully controllable in the laboratory. Compared to traditional antibody development approaches, RAMIHM is faster than regular immunization, and generates fully human mAbs without the need for humanization from traditional animal immunization. Thus, the resulted mAbs developed by RAMIHM is fully human and ready for downstream IND-enabling and/or translational studies.

In this study, we identified 3 potent and specific anti-Omicron neutralizing mAbs from Ig humanized mice by RAMIHM. Among those mAbs, MB.08 showed the high binding capacity (K_*D*_ = 7nM) and strong neutralizing ability against pseudotyped SARS-CoV-2 Omicron RBD (IC50 = 44ng/mL). All three clones are more potent than the majority of currently approved or authorized clinical RBD-targeting mAbs. Results of epitope binning experiment suggested that MB.08 might bind to sites in Omicron spike RBD with overlapping epitope(s) to PC.03 and MB.02. Nevertheless, an antibody cocktail combining MB.08 with MB.02 exhibited enhanced SARS-CoV-2 Omicron neutralization potency (IC50 = 30 ng/mL) compared to individual clones. These antibodies or their cocktail combinations are worthy of further development, such as downstream IND-enabling and/or translational studies. In general, RAMIHM can also serve as a versatile platform broadly applicable in antibody discovery against emerging pathogens or other therapeutic targets.

## Acknowledgments

We thank various members from our labs for discussions and support. We thank staffs from various Yale core facilities (Keck, YCGA, HPC, YARC, CBDS and others) for technical support. We thank Drs. Tsemperouli, Karatekin, Lin and others for providing equipment and related support. We thank various support from Department of Genetics; Institutes of Systems Biology and Cancer Biology; Dean’s Office of Yale School of Medicine and the Office of Vice Provost for Research.

## Funding

This work is supported by DoD PRMRP IIAR (W81XWH-21-1-0019) and discretionary funds to SC. The TEM core is supported by NIH grant GM132114. YCGA is supported by an NIH instrument grant 1S10OD028669-01.

## Institutional Approval

This study has received institutional regulatory approval. All recombinant DNA (rDNA) and biosafety work were performed under the guidelines of Yale Environment, Health and Safety (EHS) Committee with approved protocols (Chen 15-45, 18-45, 20-18, 20-26). All animal work was performed under the guidelines of Yale University Institutional Animal Care and Use Committee (IACUC) with approved protocols (Chen-2020-20358; Chen 2021-20068).

## Methods

### Rapid mRNA immunization of humanized mice

The full-length Omicron spike sequence used in mRNA immunization was based on two North America patients identified on Nov23^rd^, 2021. The LNP-mRNA was generated as previously described (Fang et al., 2022). Humanized mice with human IgG and IgK transgene knock-ins (ATX-GK, Alloy Therapeutics) were used for rapid mRNA immunization, according to an accelerated (two-week) vaccination schedule. Pre-immune sera were collected from the mice prior to the initiation of immunization. The mice were primed with intramuscular injection of 10μg Omicron LNP-mRNA and boosted on days 2, 4, 7 with the same dose as prime. On day 11, three days prior to sacrifice, mice received a final boost with 20μg Omicron LNP-mRNA. All mice were retro-orbital bled on days 7, 14 and anti-plasma titers were evaluated using an immunoassay as described below.

### ELISA analysis for plasma and mAbs supernatant binding to Omicron RBD protein

Plasma was extracted from surface layer by using SepMate-15 tubes with Lymphoprep gradient medium (StemCell Technologies) after centrifugation at 1200g for 20 minutes. Afterwards, antibody titers in plasma against Omicron RBD were evaluated using a direct coating ELISA. 384-well microtiter plate (Corning) were coated with 3μg/ml of Omicron RBD recombinant protein (Sino Biological 40592-V08H121) in PBS at 4°C for overnight. Plate was washed with standard wash buffer PBS-T (PBS containing 0.05% Tween 20) and blocked with blocking buffer (PBS containing 0.5% BSA) for 1 hour at room temperature (RT). Either serially diluted plasma samples or mAbs supernatant were added to plate and incubated for 1hour at RT. Wells were then washed and incubated with secondary goat anti-mouse IgG labeled with HRP (Fisher, Cat# A-10677) at 1:2500 dilution in a blocking buffer for 1h at RT. Thereafter, wells were developed using TMB substrate (Biolegend, 421101) according to the manufacturer’s protocol. The reactions were terminated with 1M H3PO4 after 20 minutes incubation at RT and optical density (OD) was measured by a spectrophotometer at 450nm (PerkinElmer EnVision 2105).

### Humanized mice B cell isolation and purification

Three sets of single B cells were collected: PBMC sample, Omicron RBD-specific memory B cell sample and CD138^+^ plasma B cell sample. PBMC cells were isolated from fresh whole blood by using SepMate-15 tubes with Lymphoprep gradient medium (StemCell Technologies) after centrifugation at 1200g for 20 minutes. Poured top layer solution that contained PBMCs from SepMate tubes to a new falcon tube and washed once with PBS+2%FBS, resuspended with PBS and stored on ice until use.

Omicron RBD-specific memory B cells were isolated from pre-enriched memory B cells by magnetic positive selection according to the manufacturer’s protocol (Miltenyi Biotec, 130-095-838). Briefly, spleen and lymph nodes were gently homogenized and red blood cells were lysed in ACK lysis buffer (Lonza). The remaining cells were washed by PBS with 2%FBS and filtered through with a 50ml falcon tube. Thereafter, memory B cells were labeled with memory B cell biotin-antibody cocktail combined with anti-biotin microbeads and isolated using a magnetic rack. Enriched memory B cells were eluted and mixed with 25ug of Omicron RBD recombinant protein with his tag and incubated for 30mins on ice. After incubation, the complex was washed and respectively incubated with anti-his-APC antibody and anti-APC microbeads. The final antigen-enrichment B cells were eluted in PBS and stored on ice until use.

Plasma B cells were collected by fragmenting and rinsing bone marrows with PBS containing 2% FBS. Non-plasma cells were labeled with a biotin-conjugated antibody cocktail combined with anti-biotin microbeads and separated using a magnetic rack according to the manufacturer’s protocol (Miltenyi Biotec, 130-092-530). Purified plasma B cells were eluted and sequentially incubated with CD138 microbeads for an additional 15 minutes at 4°C. The final CD138^+^ plasma B cells were eluted in PBS and stored on ice until use.

### Single cell VDJ sequencing and data analysis

10,000 of cells per each above collection were loaded on Chromium Next GEM Chip K Single Cell Kit. Single-cell lysis and cDNA first strand synthesis were performed using Chromium Next GEM Single Cell 5’ Kit v2 according to the manufacturer’s protocol. The barcoded single strand cDNA was isolated via a Dynabeads MyOne SILANE bead cleanup mixture. The cDNA was amplified by 14 PCR cycles and purified via SPRI bead cleanup (X0.6) according to the manufacturer’s protocol. For BCR repertoire libraries, 2 μL of amplified cDNA underwent two rounds of Target Enrichment using nested custom primer pairs specific for BCR constant regions. The target’s enrichments for heavy chain and light chain were performed in separate reactions. After each PCR reaction, the PCR products were subjected to double-sided size selection with SPRI bead cleanup (X0.6 followed by X0.8) The primers were designed by Alloy biotechnologies and synthesized by KECK.

25 ng of each target enrichment PCR product was combined, and used for library preparation, consisting of fragmentation, end repair, A-tailing, adaptor ligation (Library Construction Kit) and sample index PCR (Dual Index Kit TT Set A) according to the manufacturer’s instructions. The final library was profiled and quantified using the D1000 ScreenTape assay (Agilent) for TapeStation system. Libraries were sequenced by paired-end sequencing (26 × 91 bp) on an Illumina Miseq. All libraries were targeted for sequencing depth of 5,000 raw read pairs per cell.

For bioinformatic analysis, BCL data were converted to demultiplexed FASTQ files using Illumina Miseq controller and processed by using Cell Ranger v6.0.1 with default settings to align the reads to customized germline V and J gene references. The custom references were created by combining mouse constant genes along with human V(D)J genes. The consensus amino acid sequences of top-enriched clonotypes from each collection were selected by using the Loupe V(D)J Browser and cDNA sequences were synthesized for further molecular cloning and recombinant antibody expression.

### *In vitro* generation of recombinant mAbs

The cDNA of paired heavy- and light-chains from top-enriched IgG clonotypes were codon-optimized and respectively subcloned into human IgG1 expression vectors, based on Gibson assembly, to generate recombinant mAbs. mAbs were produced by transient transfection into Expi293F™ cells with equal amounts of paired heavy- and light-chain expression vectors using ExpiFectamine 293 transfection kit according to the manufacturer’s protocol (Thermo fisher). Five days post antibody expression, the secreted mAbs from cultured cells were collected and purified by affinity chromatography using rProtein A Sepharose Fast Flow beads according to the manufacturer’s instruction (Cytiva). Eluted mAbs were eventually kept in PBS for long-term storage after buffer exchange using Amicon Ultra-4 Centrifugal Filter (MilliporeSigma). The purified mAbs were examined by running SDS-PAGE and kept in -80°C for further usage.

### Omicron pseudovirus generation and neutralization assay

Omicron pseudovirus was generated by using a modified method from a previously described study. Briefly, full length Omicron spike gene was constructed into GFP encoding (pCCNanoLuc2AEGFP) human immunodeficiency vector backbone, then Omicron spike protein expression vectors were combined with HIV-1 structural corresponding plasmids and co-transfected into HEK-293T cells with PEI (1mg/ml, PEI MAX, Polyscience). Two-day post-transfection, viral supernatants were harvested, collected, filtered and aliquoted to use in assays.

Neutralization assays were performed by incubating pseudovirus with serial dilutions of mAbs. 10,000 cells/well of HEK-293T-hACE2 cells were seeded in a 96-well plate, 24 hours prior to assay. mAbs supernatant/purified mAbs were serially diluted in DMEM media with 10% FBS and incubated with an equal volume of purified Omicron pseudovirus at 37°C for 1 hour. Thereafter, the virus-antibody mixture was added triplicate onto HEK-293T-hACE2 cells and incubated at 37°C for additional 24 hours. Then, infected cells were counted and determined by evaluating GFP expression after 24 hours exposure to virus-antibody mixture using Attune NxT Acoustic Focusing Cytometer (Thermo Fisher). Half-maximal inhibitory concentration (IC50) for mAbs was calculated with a four-parameter logistic regression using GraphPad Prism (GraphPad Software Inc.).

### Antibody binding kinetics, epitope mapping by bio-layer interferometry (BLI)

Antibody binding kinetics for anti-Omicron RBD mAbs were evaluated by BLI on an Octet RED96e instrument (FortéBio) at RT. 25ng/ul of purified mAbs were captured on a AHC biosensor (Sartorius, 18-5060). The baseline was recorded for 60s in a running buffer (PBS, 0.02% Tween-20, and 0.05% BSA, pH 7.4). Followed by sensors were subjected to an association phase for 300s in wells containing Omicron RBD with his tag protein diluted in the buffer. In the dissociation phase, the sensors were immersed in the running buffer for 500s. The dissociation constants *K*_*D*_, kinetic constants *K*_*on*_ and *K*_*off*_ were calculated by FortéBio data analysis software.

For epitope mapping, two different antibodies were sequentially injected and monitored for binding activity to determine whether the two mAbs recognized separate or closely-situated epitopes by in-tandem approach on OCTET RED. Briefly, SARS-CoV-2 RBD-His recombinant protein (Sino Biological 40592-V08H121) was diluted with PBS to 20μg/mL, and was captured by anti-Penta-His (HIS1K) sensors (Sartorius, 18-5120). The primary antibody was diluted to 150nM with a running buffer in wells, and then sensors were firstly subjected to an association phase for 500s, the response value was recorded. Followed by sensors were subjected to the secondary antibody mixture, and the response value was recorded again. Competition tolerance was calculated by the percentage increase of response after the secondary antibody was added. The column indicates the primary antibody, and the row indicates secondary antibodies. Competition tolerance less than 25% indicates a high possibility of closely-situated epitope.

### ACE2 competition assay

3μg/ml of Omicron RBD recombinant protein (Sino Biological 40592-V08H121) was coated in a 384-well ELISA plate (Corning) at 4°C for overnight incubation. Plate was washed with standard wash buffer PBS-T (PBS containing 0.05% Tween 20) and blocked with a blocking buffer (PBS containing 0.5% BSA) for 1 hour at room temperature (RT). 50ng/mL his-tagged hACE2 protein and PBS were firstly added to plate and incubated for 1 hour at RT. Wells were washed and incubated with serially diluted purified mAbs were sequentially added and incubated for 1 hour at RT. Thereafter, wells were incubated with secondary goat anti-mouse IgG labeled with HRP (Fisher, Cat# A-10677) at 1:2500 dilution in blocking buffer for 1h at RT after washed. Finally, wells were developed using TMB substrate (Biolegend, 421101) according to the manufacturer’s protocol. The reactions were terminated with 1M H3PO4 after 20minutes incubation at RT and optical density (OD) was measured by a spectrophotometer at 450nm (PerkinElmer EnVision 2105).

### Standard statistics

Standard statistical methods were applied to non-high-throughput experimental data. The statistical methods are described in figure legends and/or supplementary Excel tables. The statistical significance was labeled as follows: n.s., not significant; * p < 0.05; ** p < 0.01; *** p < 0.001; **** p < 0.0001. Prism (GraphPad Software) and RStudio were used for these analyses. Additional information can be found in the supplemental excel tables.

### Schematic illustrations

Schematic illustrations were created with Affinity Designer or BioRender.

### Replication, randomization, blinding and reagent validations

#### Sample size

Sample size determination was performed according to similar work in the field. Replicate experiments have been performed for key data shown in this study.

#### Replication

Biological or technical replicate samples were randomized where appropriate. In animal experiments, mice were randomized by cage, sex and littermates.

#### Binding

Experiments were not blinded. It is unnecessary for animal immunization for antibody production to be blinded.

#### Antibodies and dilutions

Commercial antibodies used for various experiments are described in methods, with typical dilutions noted. For custom Antibodies generated in this study, dilutions were often serial titrations (i.e. a number of dilutions as specified in each figure). Commercial antibodies were validated by the vendors, and re-validated in house as appropriate. Custom antibodies were validated by specific antibody - antigen interaction assays, such as ELISA. lsotype controls were used for antibody validations. Eukaryotic cell lines: Cell lines are from various sources as described in methods. Cell lines were authenticated by original vendors, and re-validated in lab as appropriate. All cell lines tested negative for mycoplasma. No commonly misidentified lines involved.

#### Animals and other organisms

Laboratory animals: M. musculus, ATX strain (Alloy Tx).

## Data, resources and code availability

All data generated or analyzed during this study are included in this article and its supplementary information files. Specifically, source data and statistics for non-high-throughput experiments are provided in a supplementary table excel file. The ATX humanized mice are available via Alloy Therapeutics. Additional information related to this study are available from the corresponding author(s) upon reasonable request.

## Supplemental Figures

**Figure S1.**
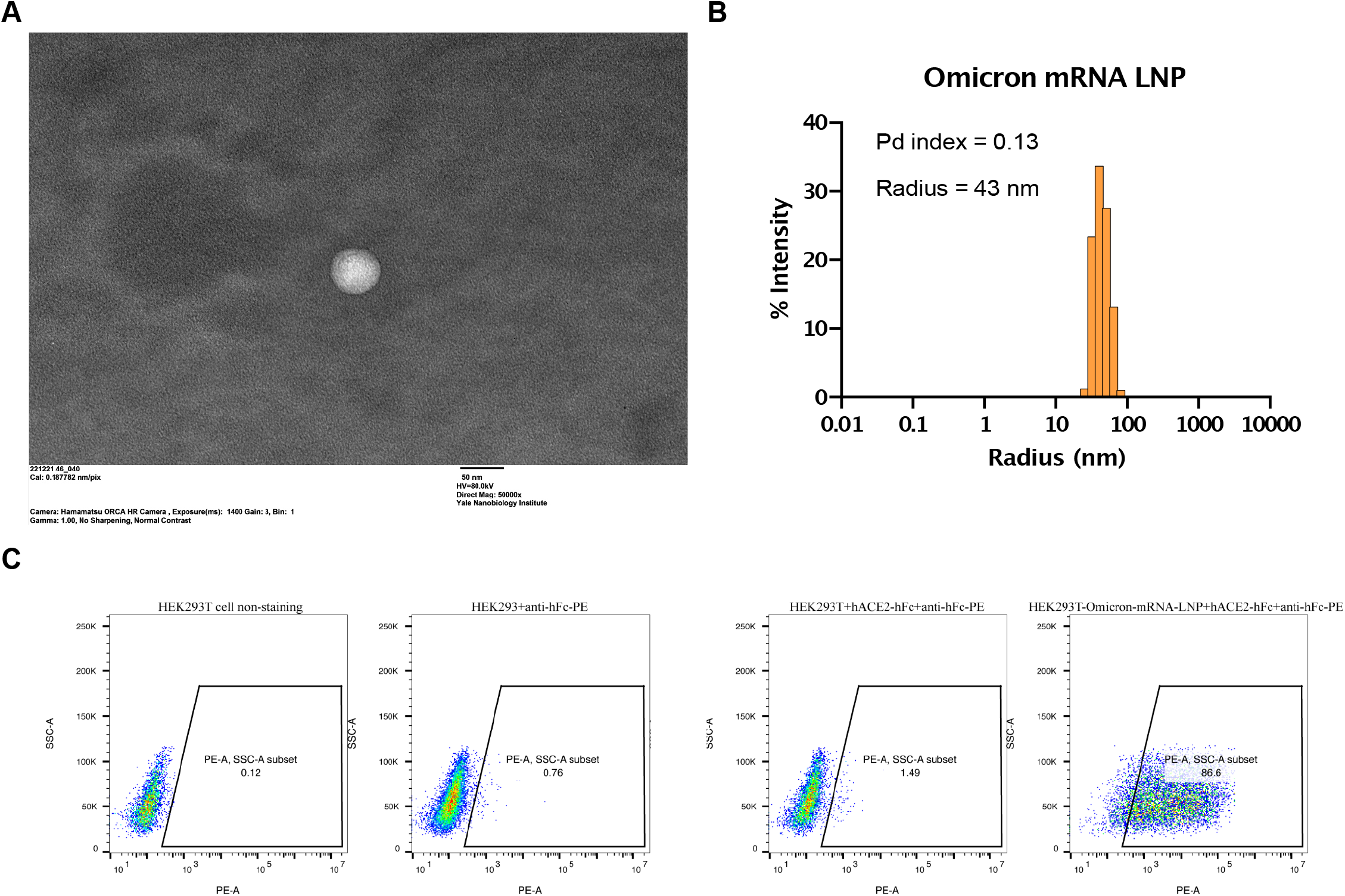
Characterization of Omicron-spike specific LNP-mRNA. **A**, Omicron LNP-mRNA image collected on transmission electron microscope. **B**, Dynamic light scattering derived histogram depicting the particle radius distribution of Omicron spike LNP-mRNA **C**, Human ACE2 receptor binding of Omicron spike expressed in 293T cells as detected by human ACE2-Fc fusion protein and PE-anti-human Fc antibody on Flow cytometry.

**Figure S2.**
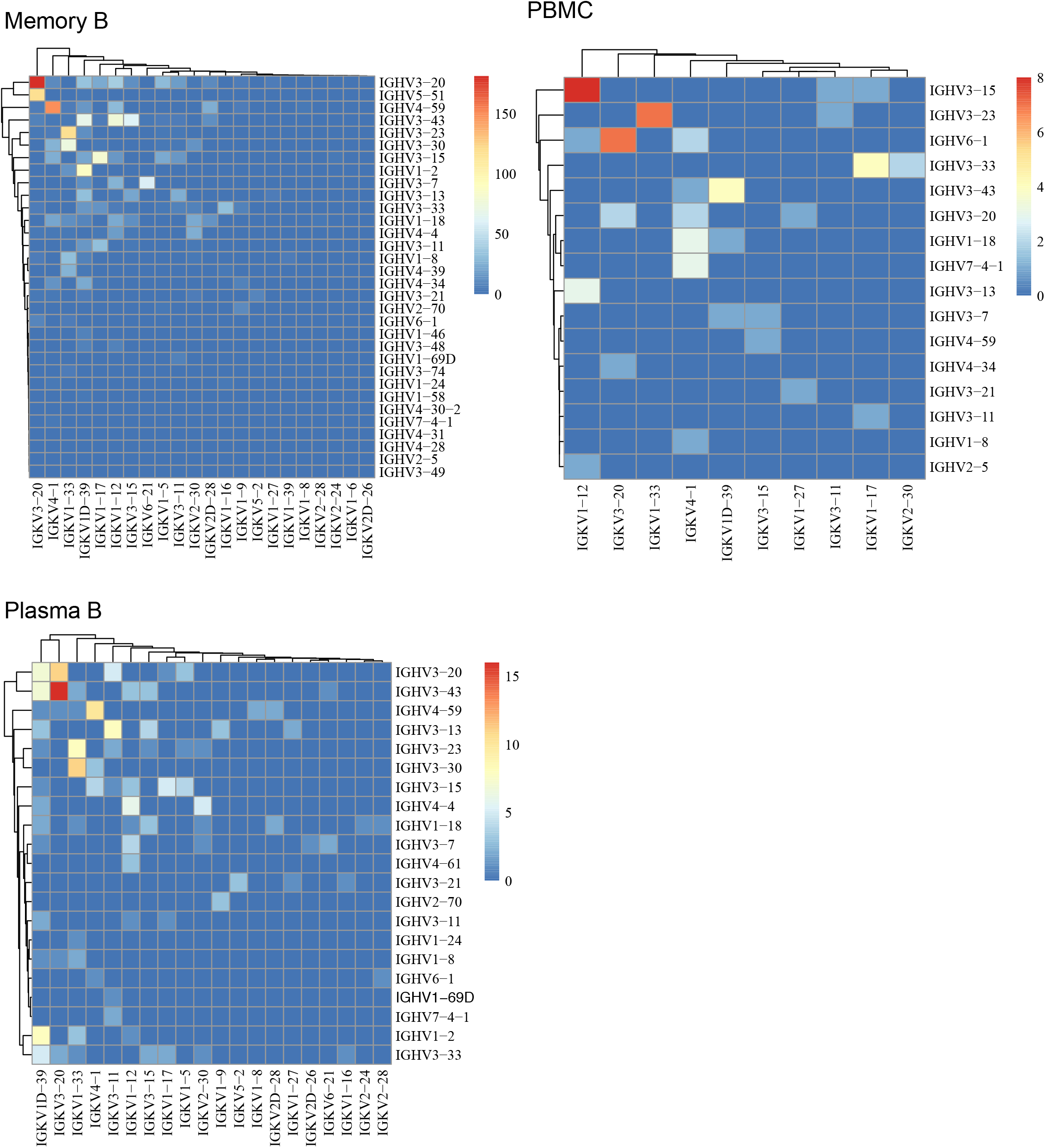
Heatmaps for non-stochastic paired BCR repertoire. Heatmaps showing the paired of immunoglobin heavy chains and light chains gene variable region segment of clonotypes in Omicron-RAMIHM mice. The reader color means the higher usage of specific VH-VL gene pairs. Memory B library, Plasma B library and PBMC library were shown in separate plots. Source data and additional statistics for experiments are in supplemental excel file(s).

**Figure S3.**
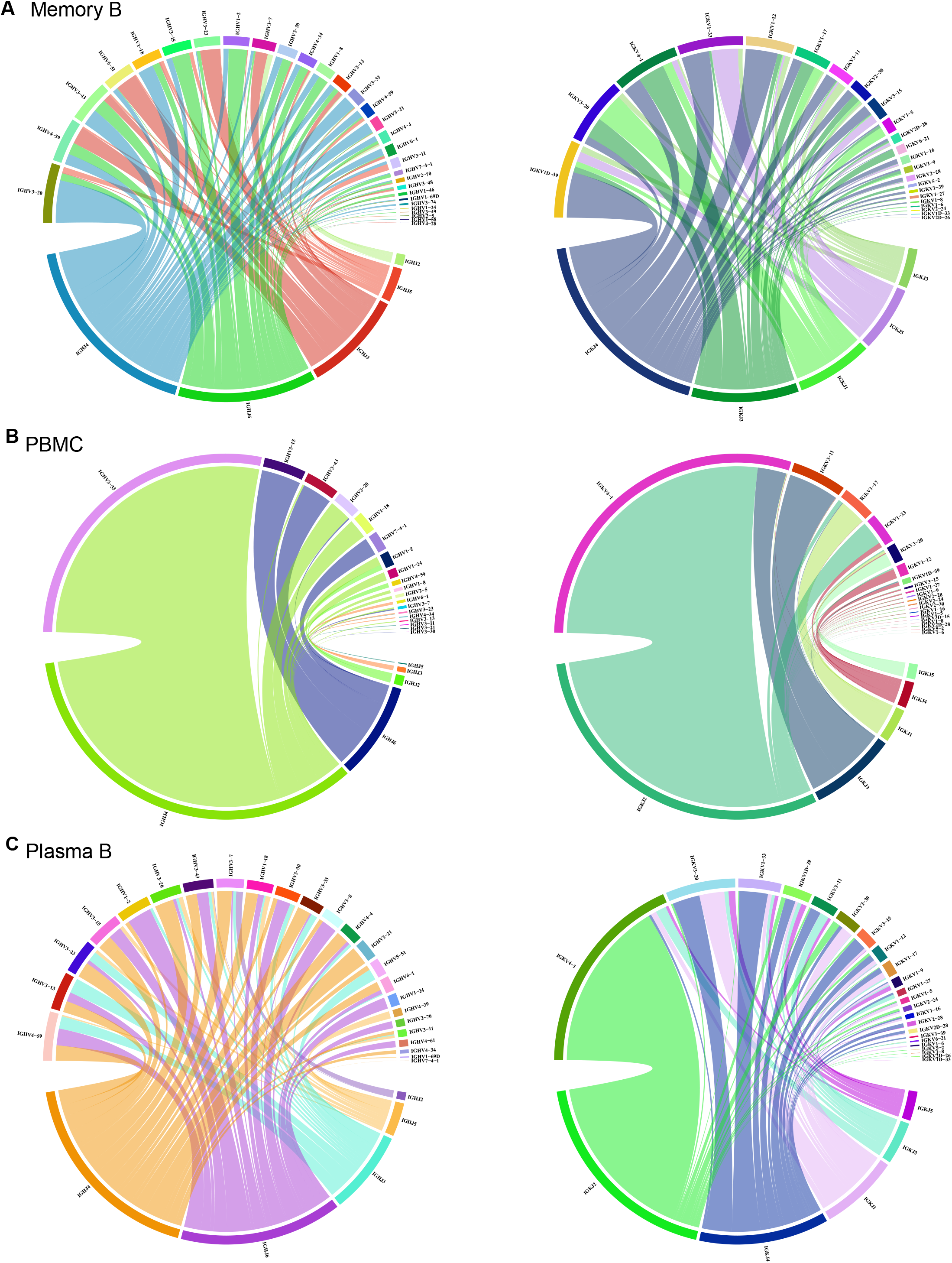
Distribution of heavy- and light-chain V/J segment recombination. Chord diagrams (circos plots) showing the distribution of all heavy- and light-chain V and J gene-segment recombination obtained in each representative library. Interconnecting lines indicate the relationship between antibodies that share V and J gene-segment at both IGH and IGL. **A**, Memory B library, **B**, PBMC library, **C**, Plasma B library. Source data and additional statistics for experiments are in supplemental excel file(s).

**Figure S4.**
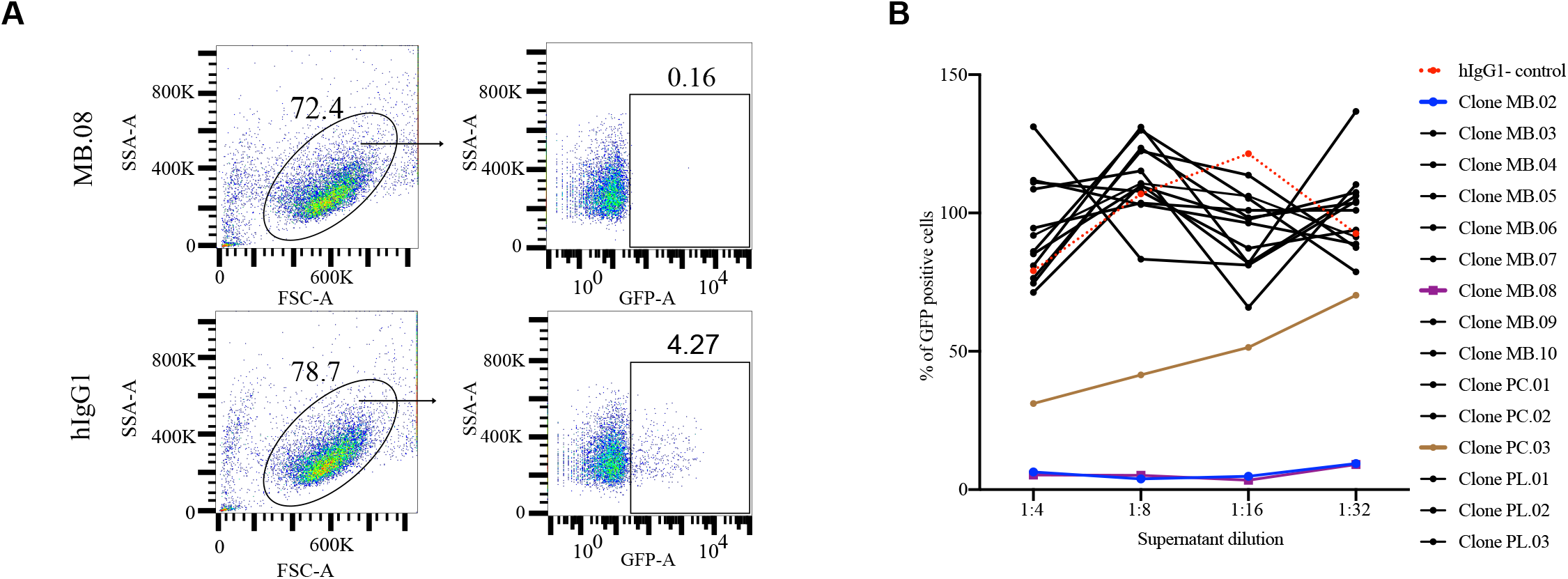
Clone screening for mAbs neutralization activity against Omicron pseudovirus. **A**, Gating strategy used for GFP-based neutralization analysis. **B**, mAbs supernatant neutralization curves in clone screening. Serial dilutions of all full length mAb clones from single BCR sequencing and control were added with Omicron pseudovirus-GFP to hACE2-O/E cells, and GFP expression was monitored and measured 24 hours after infection as a readout for virus infectivity. Data are graphed as percentage neutralization relative to virus-only infection control. Source data and additional statistics for experiments are in supplemental excel file(s).

**Figure S5.**
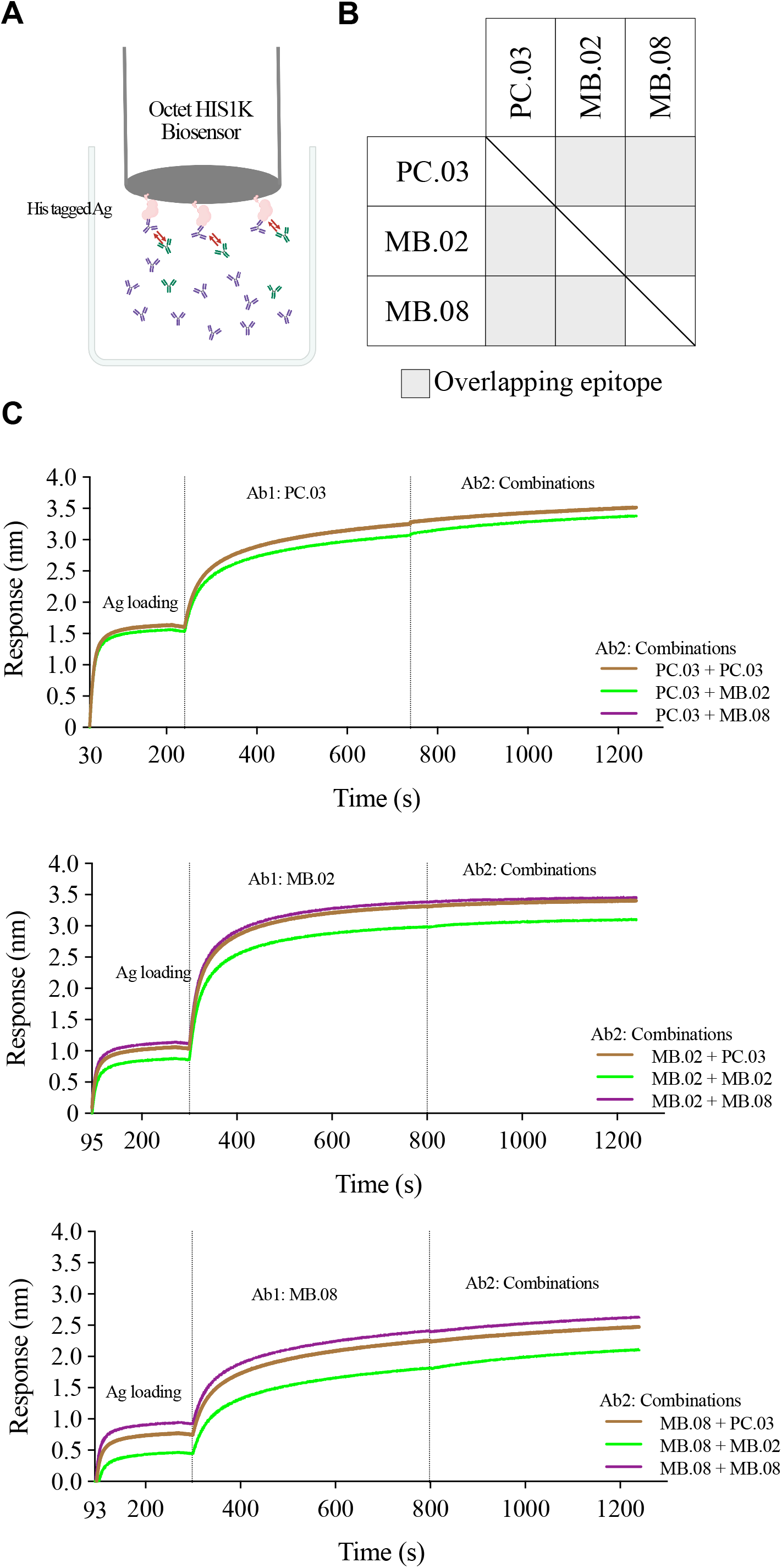
Epitope mapping through competitive binding measured by BLI. **A**, Schematic of epitope binning experiment. **B**, Summary data of BLI (C) results. The matrix presents the concluded epitope specificity for each competition experiments. The column indicated the primary loading antibody, and the row indicated the secondary antibody combinations. **C**, Epitope binning of the three potent neutralizing mAbs. Sensorgram show distinct binding patterns when pairs of testing antibodies were sequentially applied to the recombinant SARS-CoV2 Omicron RBD covalently immobilized onto a HIS1K sensor. The level of increment in response unit comparing with or without prior antibody incubation is the key criteria for determining the two mAbs recognize the separate or closely situated epitopes. Source data and additional statistics for experiments are in supplemental excel file(s).

**Figure S6.**
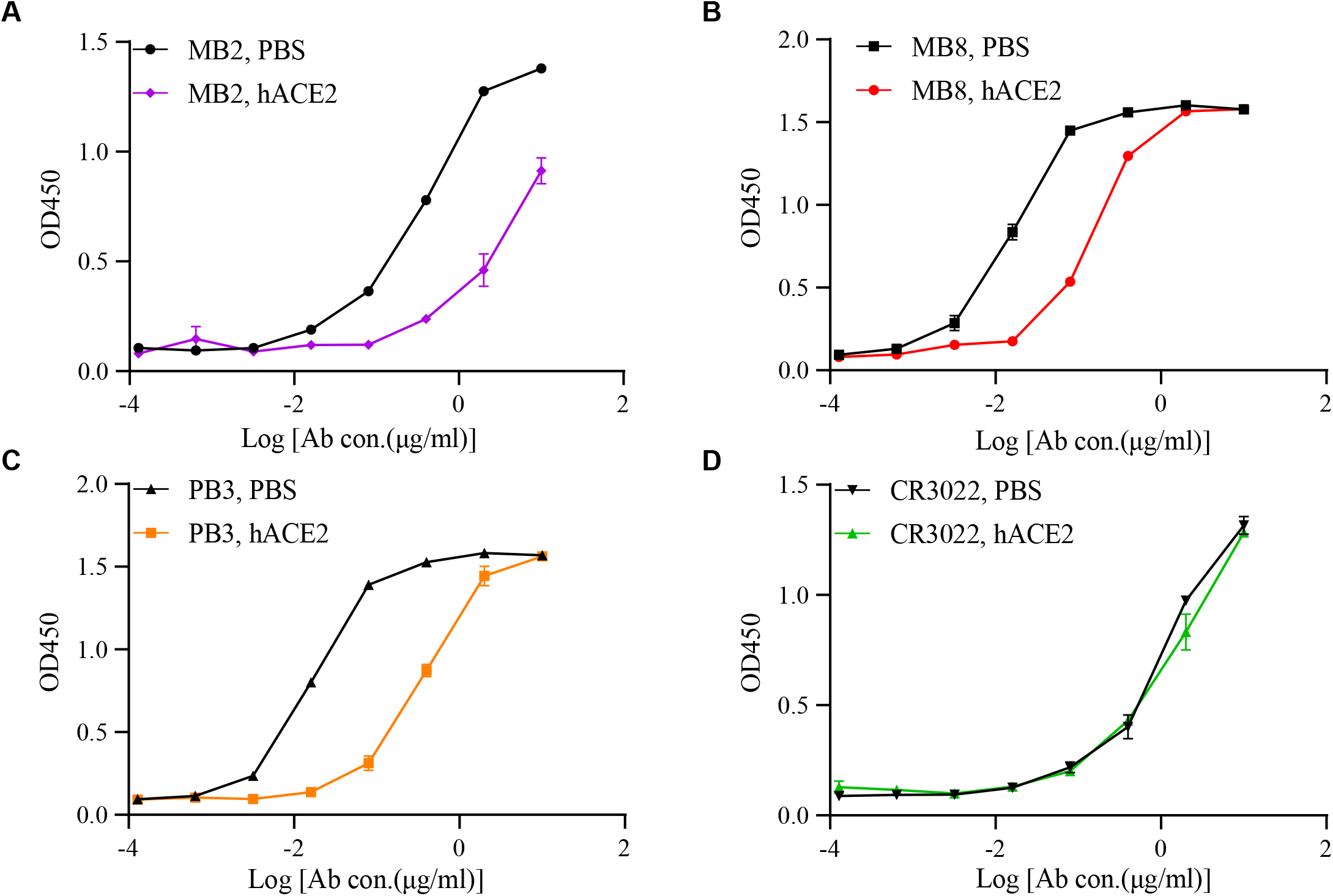
ACE2 competition for binding to SARS-CoV-2 Omicron RBD measured by ELISA. **A-D**, Curves show distinct binding patterns of ACE2 to SARS-CoV-2 Omicron RBD with or without prior antibody incubation with each testing antibody. The competition capacity of each antibody is indicated by the level of reduction in response unit of ACE comparing with or without prior antibody incubation. A commercial mAb CR3022 that binds to conserved region of spike was used as a control. **A**, MB.02 clone **B**, MB.08 clone **C**, PB.03 clone **D**, CR3022 control mAb Source data and additional statistics for experiments are in supplemental excel file(s).

**Figure S7.**
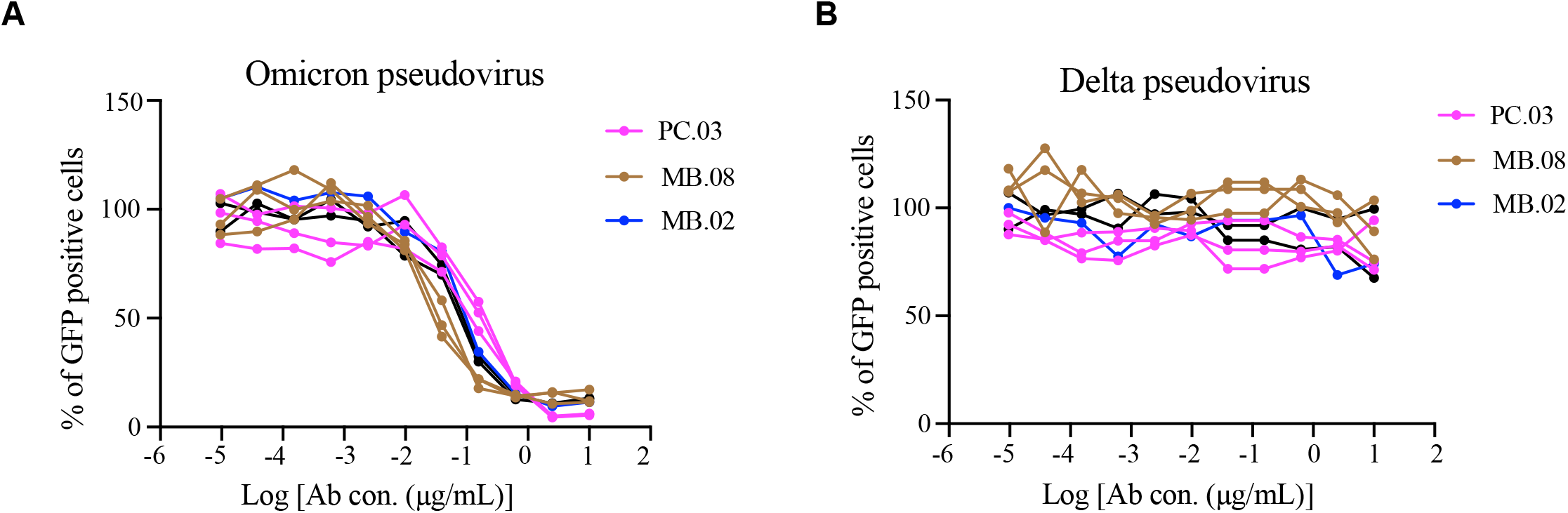
Neutralization assay of leading Omicron mAbs with pseudotyped SARS-CoV-2 variants. **A**, Individual neutralization curves for leading Omicron mAbs against Omicron pseudovirus. **B**, Individual neutralization curves for leading Omicron mAbs against Delta pseudovirus. Source data and additional statistics for experiments are in supplemental excel file(s).

## Supplemental Tables

**Key resources table (KRT)**

**Table S1. Summary of mAb efficacy against Omicron variant Source data and statistics**

Source data and statistics provided in an excel file

## Supplemental Datasets

**Dataset S1. Single cell BCR-seq processed results**

## Notes

### Competing Interest Statement

The authors have declared no competing interest.

## References

Baden, L.R., El Sahly, H.M., Essink, B., Kotloff, K., Frey, S., Novak, R., Diemert, D., Spector, S.A., Rouphael, N., Creech, C.B., et al. (2021). Efficacy and Safety of the mRNA-1273 SARS-CoV-2 Vaccine. N Engl J Med 384, 403–416.

Callaway, E. (2021). Omicron likely to weaken COVID vaccine protection. Nature 600, 367–368.

Cameroni, E., Saliba, C., Bowen, J.E., Rosen, L.E., Culap, K., Pinto, D., VanBlargan, L.A., De Marco, A., Zepeda, S.K., Iulio, J.D., et al. (2021). Broadly neutralizing antibodies overcome SARS-CoV-2 Omicron antigenic shift. bioRxiv.

Cao, Y., Wang, J., Jian, F., Xiao, T., Song, W., Yisimayi, A., Huang, W., Li, Q., Wang, P., An, R., et al. (2021). Omicron escapes the majority of existing SARS-CoV-2 neutralizing antibodies. Nature.

Carreno, J.M., Alshammary, H., Tcheou, J., Singh, G., Raskin, A., Kawabata, H., Sominsky, L., Clark, J., Adelsberg, D.C., Bielak, D., et al. (2021). Activity of convalescent and vaccine serum against SARS-CoV-2 Omicron. Nature.

Cele, S., Jackson, L., Khan, K., Khoury, D.S., Moyo-Gwete, T., Tegally, H., Scheepers, C., Amoako, D., Karim, F., Bernstein, M., et al. (2021a). SARS-CoV-2 Omicron has extensive but incomplete escape of Pfizer BNT162b2 elicited neutralization and requires ACE2 for infection. medRxiv.

Cele, S., Jackson, L., Khoury, D.S., Khan, K., Moyo-Gwete, T., Tegally, H., San, J.E., Cromer, D., Scheepers, C., Amoako, D.G., et al. (2021b). Omicron extensively but incompletely escapes Pfizer BNT162b2 neutralization. Nature.

Cerutti, G., Guo, Y., Zhou, T., Gorman, J., Lee, M., Rapp, M., Reddem, E.R., Yu, J., Bahna, F., Bimela, J., et al. (2021). Potent SARS-CoV-2 neutralizing antibodies directed against spike N-terminal domain target a single supersite. Cell Host Microbe 29, 819–833 e817.

Corbett, K.S., Flynn, B., Foulds, K.E., Francica, J.R., Boyoglu-Barnum, S., Werner, A.P., Flach, B., O’Connell, S., Bock, K.W., Minai, M., et al. (2020). Evaluation of the mRNA-1273 Vaccine against SARS-CoV-2 in Nonhuman Primates. N Engl J Med 383, 1544–1555.

Dejnirattisai, W., Huo, J., Zhou, D., Zahradnik, J., Supasa, P., Liu, C., Duyvesteyn, H.M.E., Ginn, H.M., Mentzer, A.J., Tuekprakhon, A., et al. (2022a). SARS-CoV-2 Omicron-B.1.1.529 leads to widespread escape from neutralizing antibody responses. Cell.

Dejnirattisai, W., Shaw, R.H., Supasa, P., Liu, C., Stuart, A.S., Pollard, A.J., Liu, X., Lambe, T., Crook, D., Stuart, D.I., et al. (2022b). Reduced neutralisation of SARS-CoV-2 omicron B.1.1.529 variant by post-immunisation serum. Lancet 399, 234–236.

Dhar, M.S., Marwal, R., Vs, R., Ponnusamy, K., Jolly, B., Bhoyar, R.C., Sardana, V., Naushin, S., Rophina, M., Mellan, T.A., et al. (2021). Genomic characterization and epidemiology of an emerging SARS-CoV-2 variant in Delhi, India. Science 374, 995–999.

Fang, Z., Peng, L., Lin, Q., Ren, P., KSuzuki, K., Xiong, Q., Clark, P., Lin, C., and Chen, S. (2022). SARS-CoV-2 Omicron-specific mRNA vaccine induces potent and broad antibody responses in vivo. bioRxiv.

Faria, N.R., Mellan, T.A., Whittaker, C., Claro, I.M., Candido, D.D.S., Mishra, S., Crispim, M.A.E., Sales, F.C.S., Hawryluk, I., McCrone, J.T., et al. (2021). Genomics and epidemiology of the P.1 SARS-CoV-2 lineage in Manaus, Brazil. Science 372, 815–821.

Flemming, A. (2022). Omicron, the great escape artist. Nat Rev Immunol.

Han, P., Li, L., Liu, S., Wang, Q., Zhang, D., Xu, Z., Han, P., Li, X., Peng, Q., Su, C., et al. (2022). Receptor binding and complex structures of human ACE2 to spike RBD from omicron and delta SARS-CoV-2. Cell.

Hoffmann, M., Kruger, N., Schulz, S., Cossmann, A., Rocha, C., Kempf, A., Nehlmeier, I., Graichen, L., Moldenhauer, A.S., Winkler, M.S., et al. (2022). The Omicron variant is highly resistant against antibody-mediated neutralization: Implications for control of the COVID-19 pandemic. Cell 185, 447–456 e411.

Hu, J., Peng, P., Cao, X., Wu, K., Chen, J., Wang, K., Tang, N., and Huang, A.L. (2022). Increased immune escape of the new SARS-CoV-2 variant of concern Omicron. Cell Mol Immunol 19, 293–295.

Jackson, L.A., Anderson, E.J., Rouphael, N.G., Roberts, P.C., Makhene, M., Coler, R.N., McCullough, M.P., Chappell, J.D., Denison, M.R., Stevens, L.J., et al. (2020). An mRNA Vaccine against SARS-CoV-2 - Preliminary Report. N Engl J Med 383, 1920–1931.

Johnson, B.A., Xie, X., Bailey, A.L., Kalveram, B., Lokugamage, K.G., Muruato, A., Zou, J., Zhang, X., Juelich, T., Smith, J.K., et al. (2021). Loss of furin cleavage site attenuates SARS-CoV-2 pathogenesis. Nature 591, 293–299.

Ju, B., Zhang, Q., Ge, J., Wang, R., Sun, J., Ge, X., Yu, J., Shan, S., Zhou, B., Song, S., et al. (2020). Human neutralizing antibodies elicited by SARS-CoV-2 infection. Nature 584, 115–119.

Karim, S.S.A., and Karim, Q.A. (2021). Omicron SARS-CoV-2 variant: a new chapter in the COVID-19 pandemic. Lancet 398, 2126–2128.

Krammer, F. (2020). SARS-CoV-2 vaccines in development. Nature 586, 516–527.

Liu, L., Iketani, S., Guo, Y., Chan, J.F., Wang, M., Liu, L., Luo, Y., Chu, H., Huang, Y., Nair, M.S., et al. (2021). Striking Antibody Evasion Manifested by the Omicron Variant of SARS-CoV-2. Nature.

Lopez Bernal, J., Andrews, N., Gower, C., Gallagher, E., Simmons, R., Thelwall, S., Stowe, J., Tessier, E., Groves, N., Dabrera, G., et al. (2021). Effectiveness of Covid-19 Vaccines against the B.1.617.2 (Delta) Variant. N Engl J Med 385, 585–594.

Lu, R., Zhao, X., Li, J., Niu, P., Yang, B., Wu, H., Wang, W., Song, H., Huang, B., Zhu, N., et al. (2020). Genomic characterisation and epidemiology of 2019 novel coronavirus: implications for virus origins and receptor binding. Lancet 395, 565–574.

McCallum, M., Czudnochowski, N., Rosen, L.E., Zepeda, S.K., Bowen, J.E., Walls, A.C., Hauser, K., Joshi, A., Stewart, C., Dillen, J.R., et al. (2022). Structural basis of SARS-CoV-2 Omicron immune evasion and receptor engagement. Science, eabn8652.

McCallum, M., De Marco, A., Lempp, F.A., Tortorici, M.A., Pinto, D., Walls, A.C., Beltramello, M., Chen, A., Liu, Z., Zatta, F., et al. (2021). N-terminal domain antigenic mapping reveals a site of vulnerability for SARS-CoV-2. Cell 184, 2332–2347 e2316.

Meo, S.A., Meo, A.S., Al-Jassir, F.F., and Klonoff, D.C. (2021). Omicron SARS-CoV-2 new variant: global prevalence and biological and clinical characteristics. Eur Rev Med Pharmacol Sci 25, 8012–8018.

Naranbhai, V., Garcia-Beltran, W.F., Chang, C.C., Mairena, C.B., Thierauf, J.C., Kirkpatrick, G., Onozato, M.L., Cheng, J., St Denis, K.J., Lam, E.C., et al. (2021). Comparative immunogenicity and effectiveness of mRNA-1273, BNT162b2 and Ad26.COV2.S COVID-19 vaccines. J Infect Dis.

Peng, L., Hu, Y., Mankowski, M.C., Ren, P., Chen, R.E., Wei, J., Zhao, M., Li, T., Tripler, T., Ye, L., et al. (2021). Monospecific and bispecific monoclonal SARS-CoV-2 neutralizing antibodies that maintain potency against B.1.617. bioRxiv.

Piccoli, L., Park, Y.J., Tortorici, M.A., Czudnochowski, N., Walls, A.C., Beltramello, M., Silacci-Fregni, C., Pinto, D., Rosen, L.E., Bowen, J.E., et al. (2020). Mapping Neutralizing and Immunodominant Sites on the SARS-CoV-2 Spike Receptor-Binding Domain by Structure-Guided High-Resolution Serology. Cell 183, 1024–1042 e1021.

Planas, D., Saunders, N., Maes, P., Guivel-Benhassine, F., Planchais, C., Buchrieser, J., Bolland, W.H., Porrot, F., Staropoli, I., Lemoine, F., et al. (2021). Considerable escape of SARS-CoV-2 Omicron to antibody neutralization. Nature.

Polack, F.P., Thomas, S.J., Kitchin, N., Absalon, J., Gurtman, A., Lockhart, S., Perez, J.L., Perez Marc, G., Moreira, E.D., Zerbini, C., et al. (2020). Safety and Efficacy of the BNT162b2 mRNA Covid-19 Vaccine. N Engl J Med 383, 2603–2615.

Rossler, A., Riepler, L., Bante, D., von Laer, D., and Kimpel, J. (2022). SARS-CoV-2 Omicron Variant Neutralization in Serum from Vaccinated and Convalescent Persons. N Engl J Med.

Sadoff, J., Gray, G., Vandebosch, A., Cardenas, V., Shukarev, G., Grinsztejn, B., Goepfert, P.A., Truyers, C., Fennema, H., Spiessens, B., et al. (2021). Safety and Efficacy of Single-Dose Ad26.COV2.S Vaccine against Covid-19. N Engl J Med 384, 2187–2201.

Scott, L., Hsiao, N.Y., Moyo, S., Singh, L., Tegally, H., Dor, G., Maes, P., Pybus, O.G., Kraemer, M.U.G., Semenova, E., et al. (2021). Track Omicron’s spread with molecular data. Science 374, 1454–1455.

Starr, T.N., Greaney, A.J., Dingens, A.S., and Bloom, J.D. (2021). Complete map of SARS-CoV-2 RBD mutations that escape the monoclonal antibody LY-CoV555 and its cocktail with LY-CoV016. Cell Rep Med 2, 100255.

Takashita, E., Kinoshita, N., Yamayoshi, S., Sakai-Tagawa, Y., Fujisaki, S., Ito, M., Iwatsuki-Horimoto, K., Chiba, S., Halfmann, P., Nagai, H., et al. (2022). Efficacy of Antibodies and Antiviral Drugs against Covid-19 Omicron Variant. N Engl J Med.

Tegally, H., Wilkinson, E., Giovanetti, M., Iranzadeh, A., Fonseca, V., Giandhari, J., Doolabh, D., Pillay, S., San, E.J., Msomi, N., et al. (2021). Detection of a SARS-CoV-2 variant of concern in South Africa. Nature 592, 438–443.

VanBlargan, L.A., Errico, J.M., Halfmann, P.J., Zost, S.J., Crowe, J.E., Jr., Purcell, L.A., Kawaoka, Y., Corti, D., Fremont, D.H., and Diamond, M.S. (2022). An infectious SARS-CoV-2 B.1.1.529 Omicron virus escapes neutralization by therapeutic monoclonal antibodies. Nat Med.

Viana, R., Moyo, S., Amoako, D.G., Tegally, H., Scheepers, C., Althaus, C.L., Anyaneji, U.J., Bester, P.A., Boni, M.F., Chand, M., et al. (2022). Rapid epidemic expansion of the SARS-CoV-2 Omicron variant in southern Africa. Nature.

Volz, E., Hill, V., McCrone, J.T., Price, A., Jorgensen, D., O’Toole, A., Southgate, J., Johnson, R., Jackson, B., Nascimento, F.F., et al. (2021). Evaluating the Effects of SARS-CoV-2 Spike Mutation D614G on Transmissibility and Pathogenicity. Cell 184, 64–75 e11.

Weinreich, D.M., Sivapalasingam, S., Norton, T., Ali, S., Gao, H., Bhore, R., Musser, B.J., Soo, Y., Rofail, D., Im, J., et al. (2021). REGN-COV2, a Neutralizing Antibody Cocktail, in Outpatients with Covid-19. N Engl J Med 384, 238–251.

Wolter, N., Jassat, W., Walaza, S., Welch, R., Moultrie, H., Groome, M., Amoako, D.G., Everatt, J., Bhiman, J.N., Scheepers, C., et al. (2022). Early assessment of the clinical severity of the SARS-CoV-2 omicron variant in South Africa: a data linkage study. Lancet 399, 437–446.

Zhou, P., Yang, X.L., Wang, X.G., Hu, B., Zhang, L., Zhang, W., Si, H.R., Zhu, Y., Li, B., Huang, C.L., et al. (2020). A pneumonia outbreak associated with a new coronavirus of probable bat origin. Nature 579, 270–273.

